## Supplemental Figures & Tables for "A jasmonate-responsive ERF transcription factor regulates steroidal glycoalkaloid biosynthesis genes in eggplant"

### Figure S1

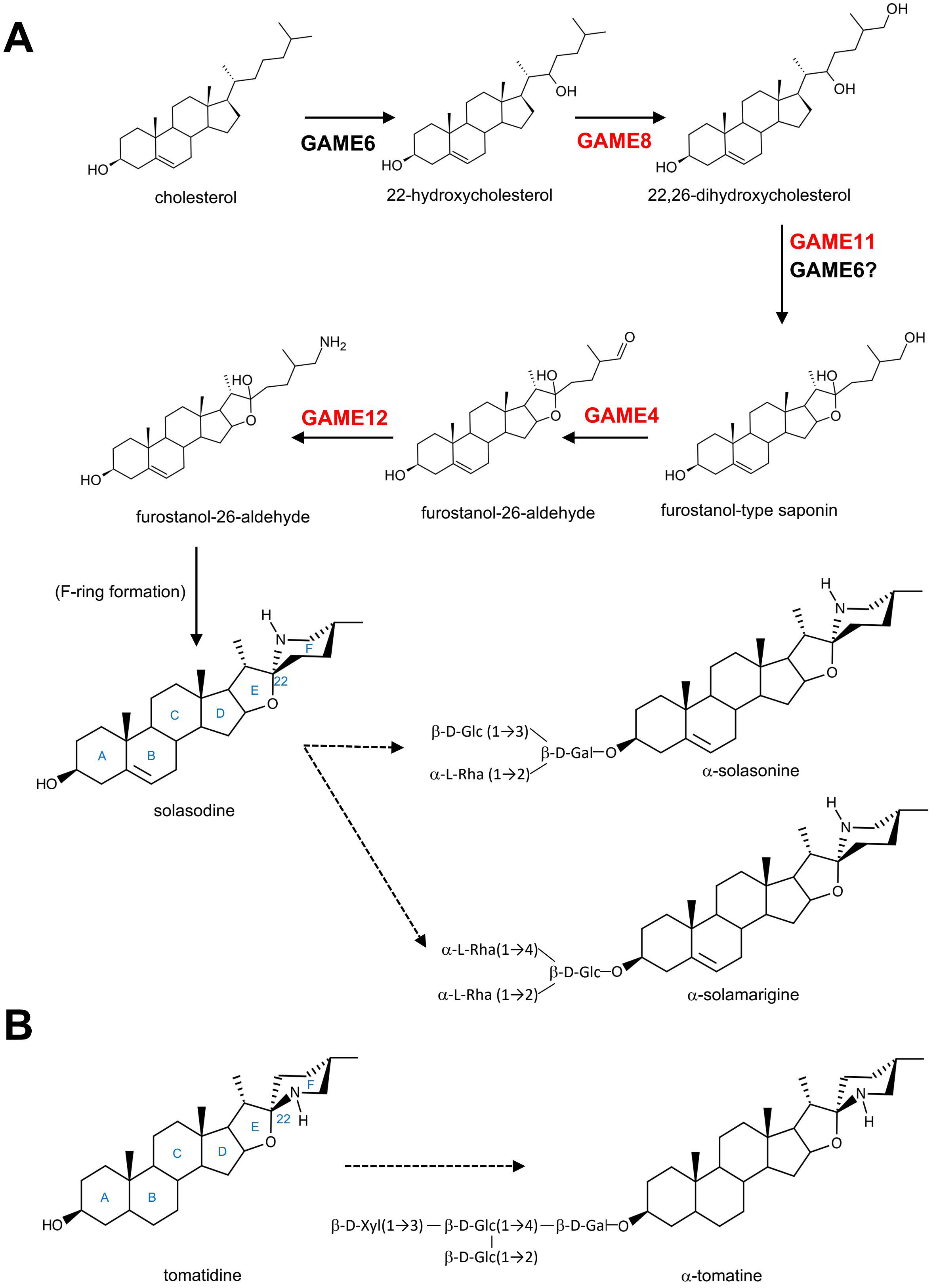


**Figure S1.** Part of the SGA biosynthesis pathway in eggplant (**A**) and tomato (**B**). Dashed arrows indicate steps involving multiple reactions. The rings are designated by letters (A to F) and C-22 are indicated in blue on the chemical structures of solasodine (**A**) and tomatidine (**B**). (**A**) Proposed reaction steps in the pathway converting cholesterol to the aglycone solasodine. Solasodine is further glycosylated, producing α-solasonine and α-solamargine, the major SGAs in eggplant. Names of metabolic enzymes deduced based on our criteria (see text and the legend for **Figure 1**) are shown in red; no gene encoding GAME6 (shown in black) was found. Hydroxylation at position 22 mediated by GAME6 in a step generating furostanol-type sapoin was presumed, but this has not been validated experimentally. (**B**) Tomatidine is glycosylated to α-tomatine the major SGA in tomato.

### Figure S2

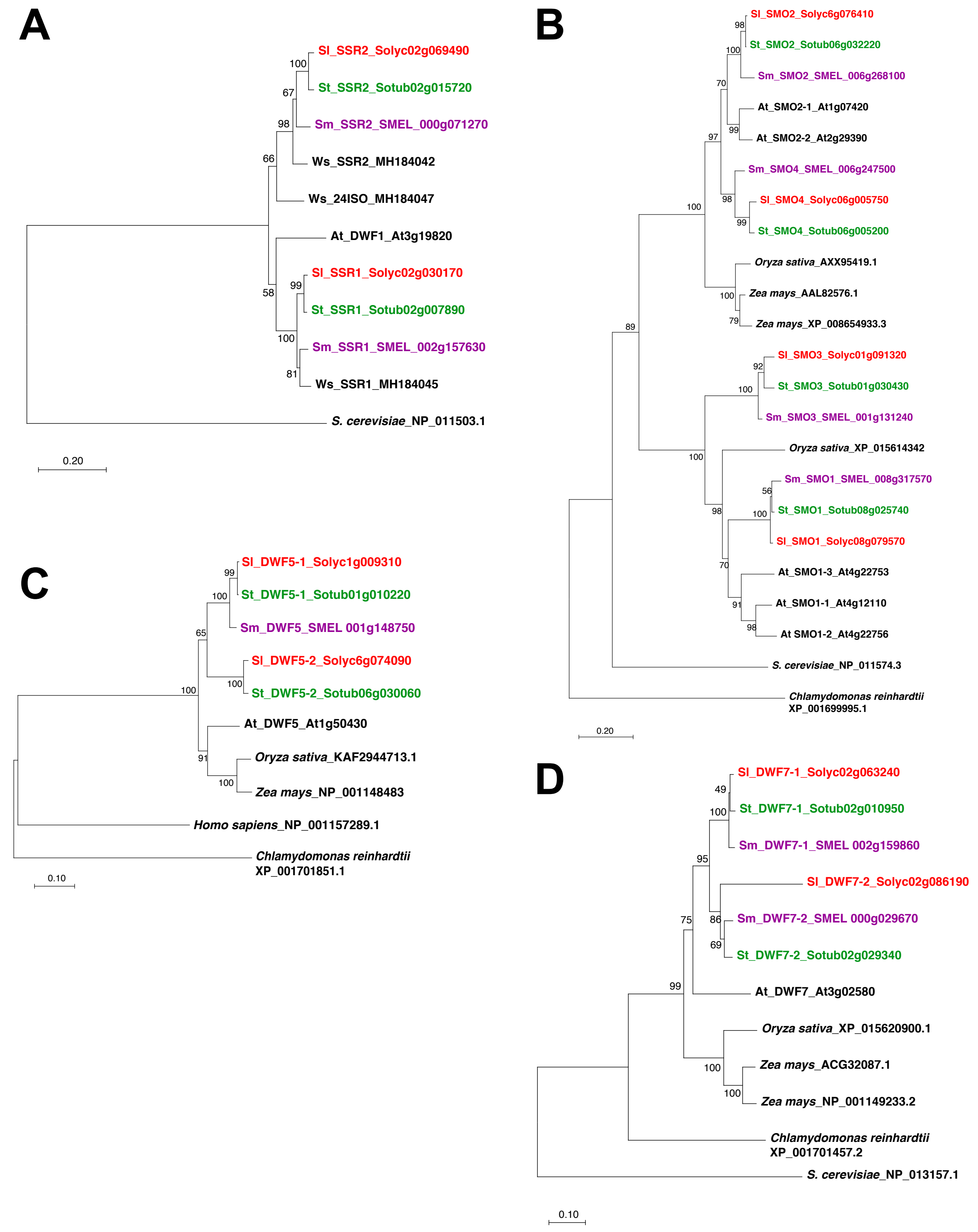


**Figure S2.** Phylogenetic relationship of proteins from eggplant and other species. Phylogenetic trees are shown for SSR (**A**), SMO (**B**), DWF5 (**C**), and DWF7 (**D**). Gene IDs or accession codes are indicated, and eggplant (purple), tomato (red), and potato (green) proteins are shown. The percentage support for 1,050 bootstrap values are indicated at branch nodes. The scale bar indicates the number of amino acid substitutions per site. Except for *SmDWF5* (SMEL001g137920), the eggplant and potato proteins were assigned to certain subgroups and named based on similarities to tomato counterparts.

### Figure S3

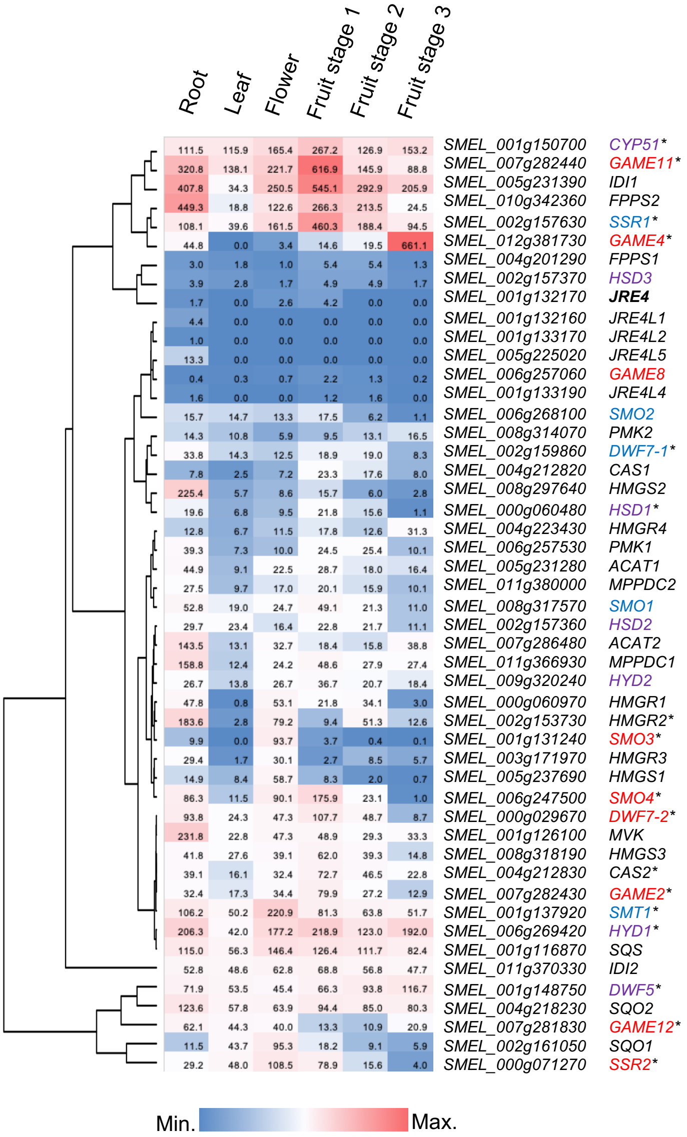


**Figure S3.** Heat map showing RPKM-normalized expression data [8] in eggplant tissues (roots, leaves, flowers, and fruits at stage 1 [8–10 d after pollination], fruit [skin and flesh] at stage 2 [pre-veraison, the onset of ripening], and fruit [skin and flesh] of stage 3 [physiological maturation]) of SGA and phytosterol biosynthesis genes. Hierarchical clustering was performed using the ward d2 method in R (http://www.R-project.org), and a heap map was drawn with Excel. The metabolic genes specific to SGA biosynthesis, phytosterol biosynthesis, and those shared by both are shown in red, blue, and purple, respectively. The genes examined here by RT-qPCR analyses are marked with asterisks. Primer sequences are listed in **Table S3.** Colors of shaded values represent RPKM values according to the bar.

### Figure S4

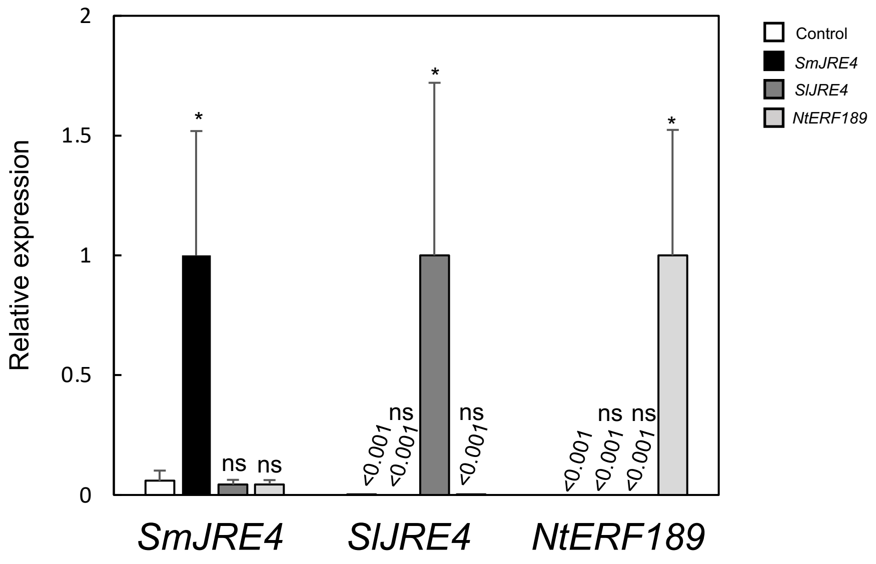


**Figure S4.** Transient overexpression of *SmJRE4*, *SlJRE4*, and *NtERF189* in eggplant leaves. The pGWB2-based vectors for overexpression of each ERF transcription factor gene were introduced into the leaves via *Agrobacterium*-mediated infection. At 2 d following the infection, transcript levels were analyzed by RT-qPCR. Primer sequences are listed in **Table S3**. The maximum values for each gene were set to 1. Detection of *SlJRE4* and *NtERF189* expression in the controls was considered to be nonspecific amplification. Significant differences relative to the controls were determined by Student's t-test. **P* < 0.01. ns; not significant.

### Figure S5

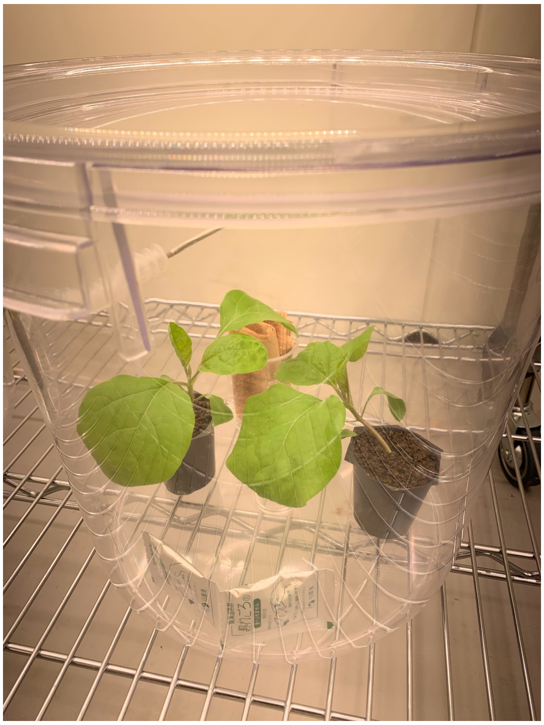

**Figure S5.** An eggplant treatment chamber. Eggplant plantlets exposed to gases of methyl JA (MJ) and ethylene in an air-tight bucket. A paper towel absorbed with 1.2 mL of MJ liquid and two packages of ethylene-generating agent Uregoro (Carto, Shizuoka, Japan) pierced with needles were placed in the bucket along with two individual plants.

**Table S1** Tomato SGA and related phytosterol enzymatic proteins, whose sequences were used as queries for BLASTP search

| name | gene ID | description |
|---|---|---|
| ACAT | Solyc05g017760 | acetyl-CoA C-acetyltransferase |
| HMGS | Solyc08g080170 | hydroxymethylglutaryl-CoA synthase |
| HMGR | Solyc02g082260 | hydroxymethylglutaryl-CoA reductase |
| MVK | Solyc01g098840 | mevalonate kinase |
| PMK | Solyc06g066310 | phosphomevalonate kinase |
| MPPDC | Solyc04g009650 | mevalonate diphosphate decarboxylase |
| IDI | Solyc04g056390 | isopentenyl diphosphate isomerase |
| FPPS | Solyc12g015860 | farnesyl diphosphate synthase |
| SQS | Solyc01g110290 | squalene synthase |
| SQO | Solyc04g077440 | squalene monooxygenase |
| CAS | Solyc04g070980 | cycloartenol synthase |
| SSR2 | Solyc02g069490 | sterol side chain reductase 2 |
| SMO3 | Solyc01g091320 | sterol methyl oxidase 3 |
| SMO4 | Solyc06g005750 | sterol methyl oxidase 4 |
| DWF7-2 | Solyc02g086180 | sterol C-5 desaturase / Dwarf 7-2 |
| DWF5-2 | Solyc06g074090 | sterol reductase / Dwarf 5-2 |
| GAME6 | Solyc07g043460 | CYP72 / Glycoalkaloid Metabolism 6 |
| GAME8 | Solyc06g061027 | CYP72 / Glycoalkaloid Metabolism 8 |
| GAME11 | Solyc07g043420 | 2-oxoglutarate-dependent dioxiganase / Glycoalksaloid Metabolism 11 |
| GAME4 | Solyc12g006460 | CYP88 / Glycoalkaloid Metabolism 4 |
| GAME12 | Solyc12g006470 | transaminase / Glycoalkaloid Metabolism 12 |
| GAME1 | Solyc07g043490 | UDP-galactosyltransferase / Glycoalkaloid Metabolism 1 |
| GAME17 | Solyc07g043480 | UDP-glucosyltransferase / Glycoalkaloid Metabolism 17 |
| GAME18 | Solyc07g043500 | UDP-glucosyltransferase / Glycoalkaloid Metabolism 18 |
| GAME2 | Solyc07g043410 | UDP-rhamnosyltransferase / Glycoalkaloid Metabolism 2 |
| HSD | Solyc02g032330 | 3β-hydroxysteroid dehydrogenase 1 |
| CPI | Solyc12g098640 | cyclopropylsterol isomerase |
| CYP51 | Solyc01g008110 | sterol C14-demethylase / CYP51 |
| HYD2 | Solyc09g009040 | Δ14-sterol reductase |
| HYD1 | Solyc06g082980 | 3β-hydroxysteroid-Δ8Δ7-isomerase |
| SMT | Solyc10g080150 | sterol methyltransferase 1 |
| SMO1 | Solyc08g079570 | sterol methyl oxidase 1 |
| SMO2 | Solyc06g076410 | sterol methyl oxidase 2 |
| DWF7-1 | Solyc02g063240 | sterol C-5 desaturase / Dwarf 7-1 |
| DWF5-1 | Solyc01g009310 | sterol reductase / Dwarf 5-1 |
| SSR1 | Solyc02g030170 | sterol side chain reductase 1 |

**Table S2** Genes involved in SGA biosynthesis and related pathways in eggplant.
Genes, whose expression was analyzed by RT-qPCR, are colored in yellow. Primers used to detect them are listed in **Table S3**.

| gene name | gene ID | description |
|---|---|---|
| *ACAT1* | SMEL_005g231280 | acetyl-CoA C-acetyltransferase 1 |
| *ACAT2* | SMEL_007g286480 | acetyl-CoA C-acetyltransferase 2 |
| *HMGS1* | SMEL_005g237690 | hydroxymethylglutaryl-CoA synthase 1 |
| *HMGS2* | SMEL_008g297640 | hydroxymethylglutaryl-CoA synthase 2 |
| *HMGS3* | SMEL_008g318190 | hydroxymethylglutaryl-CoA synthase 3 |
| *HMGR1* | SMEL_000g060970 | hydroxymethylglutaryl-CoA reductase 1 |
| *HMGR2* | SMEL_002g153730 | hydroxymethylglutaryl-CoA reductase 2 |
| *HMGR3* | SMEL_003g171970 | hydroxymethylglutaryl-CoA reductase 3 |
| *HMGR4* | SMEL_004g223430 | hydroxymethylglutaryl-CoA reductase 4 |
| *MVK* | SMEL_001g126100 | mevalonate kinase |
| *PMK1* | SMEL_006g257530 | phosphomevalonate kinase 1 |
| *PMK2* | SMEL_008g314070 | phosphomevalonate kinase 2 |
| *MPPDC1* | SMEL_011g366930 | mevalonate diphosphate decarboxylase 1 |
| *MPPDC2* | SMEL_011g380000 | mevalonate diphosphate decarboxylase 1 |
| *IDI1* | SMEL_005g231390 | isopentenyl diphosphate isomerase 1 |
| *IDI2* | SMEL_011g370330 | isopentenyl diphosphate isomerase 1 |
| *FPPS1* | SMEL_004g201290 | farnesyl diphosphate synthase 1 |
| *FPPS2* | SMEL_010g342360 | farnesyl diphosphate synthase 2 |
| *SQS* | SMEL_001g116870 | squalene synthase |
| *SQO1* | SMEL_002g161050 | squalene monooxygenase 1 |
| *SQO2* | SMEL_004g218230 | squalene monooxygenase 1 |
| *CAS1* | SMEL_004g212820 | cycloartenol synthase 1 |
| *CAS2* | SMEL_004g212830 | cycloartenol synthase 1 |
| *SSR2* | SMEL_000g071270 | sterol side chain reductase 2 |
| *SMO3* | SMEL_001g131240 | sterol methyl oxidase 3 |
| *SMO4* | SMEL_006g247500 | sterol methyl oxidase 4 |
| *DWF7-2* | SMEL_000g029670 | sterol C-5 desaturase / Dwarf 7-2 |
| *GAME8* | SMEL_006g257060 | CYP72 / Glycoalkaloid Metabolism 8 |
| *GAME11* | SMEL_007g282440 | 2-oxoglutarate-dependent dioxiganase / Glycoalksaloid Metabolism 11 |
| *GAME4* | SMEL_012g381730 | CYP88 / Glycoalkaloid Metabolism 4 |
| *GAME12* | SMEL_007g281830 | transaminase / Glycoalkaloid Metabolism 12 |
| *GAME2* | SMEL_007g282430 | UDP-rhamnosyltransferase / Glycoalkaloid Metabolism 2 |
| *HSD1* | SMEL_000g060480 | 3β-hydroxysteroid dehydrogenase 1 |
| *HSD2* | SMEL_002g157360 | 3β-hydroxysteroid dehydrogenase 2 |
| *HSD3* | SMEL_002g157370 | 3β-hydroxysteroid dehydrogenase 3 |
| *CYP51* | SMEL_001g150700 | sterol C14-demethylase / CYP51 |
| *HYD2* | SMEL_009g320240 | Δ14-sterol reductase |
| *HYD1* | SMEL_006g269420 | 3β-hydroxysteroid-Δ8Δ7-isomerase |
| *DWF5* | SMEL_001g148750 | sterol reductase / Dwarf 5 |
| *SMT1* | SMEL_001g137920 | sterol methyltransferase 1 |
| *SMO1* | SMEL_008g317570 | sterol methyl oxidase 1 |
| *SMO2* | SMEL_006g268100 | sterol methyl oxidase 2 |
| *DWF7-1* | SMEL_002g159860 | sterol C-5 desaturase / Dwarf 7-1 |
| *SSR1* | SMEL_002g157630 | sterol side chain reductase 1 |
| *JRE4* | SMEL_001g132170 | Jasmonate-responsive ERF 4 |
| *JRE4L1* | SMEL_001g132160 | JRE4-like 1 |
| *JRE4L2* | SMEL_001g133170 | JRE4-like 2 |
| *JRE4L3* | SMEL_001g133180 | JRE4-like 3 |
| *JRE4L4* | SMEL_001g133190 | JRE4-like 4 |
| *JRE4L5* | SMEL_005g225020 | JRE4-like 5 |

**Table S3** Oligonucleotide primers used in RT-qPCR analysis

| gene name | gene ID | forward/reverse (F/R) | sequence (5' to 3') |
|---|---|---|---|
| *Sm HMGR2* | SMEL_002g153730 | F | ATGGACGTTCGCCGGAGATCTG |
| | | R | CAGAGGCATCAATGAGAAGGGTGTC |
| *Sm CAS2* | SMEL_004g212830 | F | AATGTTTTGTTTTCCTTACAGAATAGTGG |
| | | R | TTGCAGCTGAGGAACACTCTACATACGG |
| *Sm SSR2* | SMEL_000g071270 | F | AGCAATATGTAAAACTTACCTACAAACCTG |
| | | R | CTACCATCTCTGGAACTTTAGAAGAATTGTC |
| *Sm SMO3* | SMEL_001g131240 | F | TTGCCATTTGGGAGCATTGAAGAG |
| | | R | GGTGTTGTGACAATATAGATAGAAATCTG |
| *Sm SMO4* | SMEL_006g247500 | F | ATGGCCTCCATGATCGAATCTGC |
| | | R | GAGTCCAGACAAGAAGAAGGCACTC |
| *Sm DWF7-2* | SMEL_000g029670 | F | AACAAGCAAAACACACTTTCCCCGTTTGCTG |
| | | R | GGAATATGAGCCCTATGTGTGTAGTGAAATGC |
| *Sm GAME11* | SMEL_007g282440 | F | GACCTCCTCTCCAACTATTCAAGC |
| | | R | TAGCTTTTCCCAAATCAATTACTGG |
| *Sm GAME4* | SMEL_012g381730 | F | AAGCCAAGCATTATTATCACAAAGCCAG |
| | | R | CTTATCTTCTTCGATTGATGTCGATAGTCC |
| *Sm GAME12* | SMEL_007g281830 | F | CAAAACCTTCTTTGGATCTTGCAAAGGAGC |
| | | R | GCATTGTTGTAATACCAGACCAGCTTCACCTGG |
| *Sm GAME2* | SMEL_007g282430 | F | ATGGCAACGGAAGTGAAGGAAC |
| | | R | GCCATGGAGGGCGAAGAGTCTAGC |
| *Sm HSD* | SMEL_002g157360 | F | TTCATAGCTTCTGGCTTCACC |
| | | R | CCATTATTCTCTGAGGAAGTCTCC |
| *Sm CYP51* | SMEL_001g150700 | F | ATGGAGTTAGGTGACAACAAGATTCTG |
| | | R | CCATGATTTGATCACTGGAGGCAAACG |
| *Sm HYD2* | SMEL_009g320240 | F | CTCCATTGACACTGCATATTAGGC |
| | | R | GAAGGAGTGAGAGAAATGAG |
| *Sm HYD1* | SMEL_006g269420 | F | ATGACAATTCAGGGAGAAGCTCAC |
| | | R | CAGGAAGGATGAGATGCCATACACAC |
| *Sm DWF5* | SMEL_001g148750 | F | ATGGCAGAGACCAAGTTGGTACAC |
| | | R | CATCAGCGTGTACATTTGTATACC |
| *Sm SMT1* | SMEL_001g137920 | F | GCACGGCGCTTTTGATCTGGC |
| | | R | CTTCTTCTTCACCTCCATAATAACC |
| *Sm DWF7-1* | SMEL_002g159860 | F | ACGACTACTTGAATTTGTTTGTCG |
| | | R | GCGGAGCCATCCTTGAAACCAATGAG |
| *Sm SSR1* | SMEL_002g157630 | F | ATGCTATACCATGGTCTCAAGGGACTCTG |
| | | R | AAGCCTGTGCAATCTCTTTCAGATTACC |
| *Sm JRE4* | SMEL_001g132170 | F | CGATTTTTCTCCAAGTATTTCGTCG |
| | | R | CTTTGTTTCCTCCGGGGGCTCACATCC |
| *Sm CYP* | SMEL_001g116150 | F | ATCCTTGTCCATGGCTAATGC |
| | | R | ATGCCCTCAACAACTTGTCC |
| *Sl JRE4* | Solyc01g090340 | F | CGATTTTTTCGAAACTCTTTCC |
| | | R | TGTTTCCTCCGGTGTTACGG |
| *Nt ERF189* | AB827951.1 | F | GCAGCTTCGACTGCAGCTTCCTC |
| | | R | CTCCTCGGACTCGGAGCACTTC |
